# The Pattern of Bacteria and its Resistance on Adult Sepsis Patient at Dr. Moewardi General Hospital, Indonesia

**DOI:** 10.1101/044032

**Authors:** Andika Dwi Mahendra, M. Kuswandi, Ika Trisharyanti D. Kusuma

## Abstract

**Background:** Sepsis incidence which related with morbidity and mortality, is globally increasing. Sepsis occurs because of severe infections. Sepsis has life-threatening potential with organ disfungction complication, septic shock, and death. Alteration of resistance pattern always change at certain period. The aim of the study is to investigate the bacteria pattern and its resistance from adult sepsis patient at Dr. Moewardi Regional General Hospital.

**Methods:** Isolated bacteria from adult sepsis patient’s blood at Dr. Moewardi Regional General Hospital. 7 bacteria isolates were taken from blood of adult sepsis patient (September - October 2014) and 46 bacteria isolates of secondary data (January - March 2014) at Dr. Moewardi Regional General Hospital. Bacteria isolation was performed based on standard Laboratory of Microbiology of Dr. Moewardi Regional General Hospital. 7 bacteria isolates were tested by using disc diffusion antibiotic on Mueller Hinton and coupled by 46 secondary data.

**Results:** The bacteria pattern from adult sepsis patients were *Staphylococcus haemolyticus* (15,09%), *Staphylococcus hominis* (15,09%), *Escherichia coli* (13,21%), and *Acinetobacter baumannii* (11,32%). Resistance pattern of bacteria on adult sepsis with level of resistant more than 50% as *Staphylococcus haemolyticus* (ciprofloxacin, erythromycin, levofloxacin, moxifloxacin, and gentamicin), *Escherichia coli* (gentamicin, ciprofloxacin, and levofloxacin), and *Acinetobacter baumannii* (gentamicin, ciprofloxacin, levofloxacin, and ceftazidime)

**Conclutions:** Based on this research, revealed that the most common bacteria caused sepsis was Gram negative about 64,64% of all bacteria on sepsis patient. The most frequent cause of sepsis was *Staphylococcus hominis, Staphylococcus haemolyticus*, and *Escherichia coli.*

## Background

Infection can occur because of infection agents such as bacteria, fungus, virus, and protozoa [1]. Bacteremia is state which presence of living bacteria on blood stream [2]. Bacteremia is state that determine to occur sepsis. Further, sepsis can develop to be sepsis shock and also renal and hepar disfunction with hypotension [3].

Bacteria is one of sepsis’s cause. It comes from previous infections such as pulmonary infection, urinary tract infection, central nerve system infection, or *Community acquired methicillin-resistant Staphylococcus aureus* [3, 4]. Sepsis patients will have possibility for developing complications such as organ complications. One of factor increasing mortality on sepsis patient was organ disfunction. Mortality of sepsis patiens were estimated about 15% and increased to be 70% if patiens suffer three or more organ disfunction including pulmonary, renal, or heart [5].

Development of bacteria resistance was increasing quickly by discovering resistant bacteria against antibiotics in 1979 – 2011. Some bacteria had been being resistant against antibiotics, for instance gentamicin-R *Enterococcus*, vancomycin-R *Enterococcus*, levofloxacin-R *Pneumococcus*, imipenem-R *Enterobacteriaceae*, vancomicin-R *Staphylococcus*, ceftriaxone-R *Nesseria gonorrhoeae*, dan ceftaroline-R *Staphylococcus* [6]. Research about *Escherichia coli* resistance showed that 21 bacteria isolates (0,6%) were resistant against ampicillin, chloramphenicol, gentamicin, ciprofloxacine, cefotaxime, and trimethoprim /sulfamethoxazole [7]. Lewis et al. reported that resistance also occured against ceftriaxone and imipenem on *Acinetobacter spp.* (28,6% dan 10%), *Pseudomonas aeruginosa* (46,7% and 3,8%), and *Enterobacter spp.* (16% and 0%) [8]. Research was performed at ICU room, Fatmawati hospital, Indonesia, revealed that *Pseudomonas aeruginosa, Staphylococcus epidermidis*, and *Escherichia coli* were resistant against meropenem, gentamicin, and levofloxacine [9].

## Method

Bacteria was isolated by standart of microbiology laboratory in Dr. Moewardi regional general hospital and then earned pure bacteria isolate. The bacteria isolates were grown on NA (Nutrient Agar) and brought to microbiology laboratory of Faculty of Pharmacy, MUS, for being performed sensitivity test against antibiotics. Sensitivity test was carried out using disc diffusion and the results were interpreted based on CLSI (*Clinical and Laboratory Standards Institute*).

The bacteria isolates, from NA, were taken by using sterile inoculation loop and put into BHI medium then shaked by shaker for 2 hours. After that, took 200 μL of BHI solution and diluted by NaCl 0,9% until having similar turbidity with Mc Farland (1,5 x 10^8^ CFU/mL). Bacteria suspension was taken 100 μL then put it on Mueller-Hinton agar and distributed it evenly. The next step, 8 antibiotic discs were put on Mueller-Hinton agar surface and incubated on 37°C for 24 hours. The inhibition zone of each antibiotic discs were measured on the next day.

Analysis of the result of bacteria sensitivity was determined by measuring inhibition zone of each antibiotic discs and compared by CLSI standart (ampicillin, gentamicin, levofloxacine, vancomycin, metronidazole, meropenem, rifampicin, and ceftriaxone), and coupled by secondary data (sensitivity test data from microbiology laboratory of Dr. Moewardi general hospital). Based on the previous result (marger primary and secondary data), so we could classify level of bacteria sensitivity on adult sepsis, included sensitive (S) or resistant (R).

This research is ethically approved by Health Research Ethics Committee Dr. Moewardi General Hospital/School of Medicine Sebelas Maret University of Surakarta. Ethical Clearance number is 476/ VII/HREC/2014.

## Result

This research was carried out to determine bacteria and its resistance pattern on adult sepsis patient against some antibiotics at Dr. Moewardi general hospital in 2014. Patient distribution based on age and gender at Dr. Moewardi regional general hopital can be seen on below (table 1). Based on table 1, sepsis frequently occurs on patients who older than 49 years old.

**Table 1.** Distribution of bacteria isolates based on age and gender at Dr. Moewardi general hospital in 2014

|  | Total | Percentage (%) |
| --- | --- | --- |
| Gender |  |  |
| Man | 27 | 58,70 |
| Woman | 19 | 41,30 |
| Total | 46 | 100,00 |
| Age (year) |  |  |
| 20-30 | 3 | 6,52 |
| 31-40 | 8 | 17,40 |
| 41-49 | 6 | 13,04 |
| >49 | 29 | 63,04 |
| Total | 46 | 100,00 |

**Table 2.** Result of bacteria isolates from adult sepsis patients at Dr. Moewardi regional general hospital in 2014

| Bacteria | Primary data | Secondary data | Total | Percentage (%) |
| --- | --- | --- | --- | --- |
| Gram-positive |  |  |  |  |
| *Staphylococcus hominis* |  | 8 | 8 | 15,09 |
| *Staphylococcus haemolyticus* | 1 | 7 | 8 | 15,09 |
| *Staphylococcus aureus* |  | 5 | 5 | 9,43 |
| *Staphylococcus epidermidis* | 1 | 4 | 5 | 9,43 |
| *Staphylococcus capitis* |  | 1 | 1 | 1,89 |
| *Enterobacter cloacae* |  | 1 | 1 | 1,89 |
| *Enterococcus faecium* |  | 1 | 1 | 1,89 |
| Total | 2 | 27 | 29 | 100,00 % |
| Gram-negative |  |  |  |  |
| *Escherichia coli* | 3 | 4 | 7 | 13,21 |
| *Acinetobacter baumannii* | 1 | 5 | 6 | 11,32 |
| *Klebsiella pneumoniae* |  | 6 | 6 | 11,32 |
| *Achromobacter xylosoxidans* |  | 2 | 2 | 3,77 |
| *Proteus mirabilis* |  | 1 | 1 | 1,89 |
| *Pseudomonas aeruginosa* | 1 |  | 1 | 1,89 |
| *Salmonella ssp* |  | 1 | 1 | 1,89 |
| Total | 7 | 46 | 53 | 100,00 % |

Total of bacteria isolates were 53 items. The sensitivity of 7 isolates from adult sepsis patients (September-October 2014) were tested using antibiotic discs by antibiotic disc diffusion method, and than coupled by 46 secondary data (January-March 2014) from microbiology laboratory of Dr. Moewardi general hospital.

Blood sample was taken from adult sepsis patient after the patient got empiric antibiotic treatments. Sampling before or after antibiotic treatment will affect on the result of bacteria pattern. Number of pathogen bacteria would be reduced by antibiotic treatment. But, it can be minimized by compounds (in growth media for blood sample) that neutralize the antibiotic effect, so the pathogen bacteria could grow well. 7 bacteria isolates consist of Gram-positive and Gram-negative bacteria covering *Staphylococcus haemolyticus*,

*Staphylococcus epidermidis, Acinetobacter baumannii, Pseudomonas aeruginosa*, and three *Escherechia coli.* The seven isolates were tested their resistance against eight antibiotics (ampicillin, gentamicin, levofloxacine, vancomycin, metronidazole, meropenem, rifampicin, and ceftriaxone). The results were shown on table 3 and table 4.

**Table 3.** The result of sensitivity test of Gram-positive from adult sepsis patient

| Bacteria<br>(bacteria code) | Antibiotics |  |  |  |  |  |  |  |  |  |  |  |  |  |  |  |
| --- | --- | --- | --- | --- | --- | --- | --- | --- | --- | --- | --- | --- | --- | --- | --- | --- |
|  | MEM<sup>1</sup> |  | CN<sup>1</sup> |  | LEV<sup>1</sup> |  | VA<sup>1</sup> |  | MTZ<sup>2</sup> |  | AMP<sup>2</sup> |  | RD<sup>2</sup> |  | CRO<sup>2</sup> |  |
|  | IZ | C | IZ | C | IZ | C | IZ | C | IZ | C | IZ | C | IZ | C | IZ | C |
| *Staphylococcus hemolyticus* (618D) | 8,5 | R | 9 | R | 14 | R | 14,5* | R | - | R | - | R | - | R | 17 | R |
| *Staphylococcus epidermidis* (68 D) | 31,7 | S | 19 | S | 28,8 | S | 13,8 | R | - | R | 23 | R | 12 | R | 17,5 | R |

<sup>1</sup>: antibiotic on antibiotic guideline for sepsis patient at Dr. Moewardi regional general hospital; <sup>2</sup>: antibiotic on prescription for sepsis patient; HZ: inhibition zone (mm)
\* = irradical; - : no inhibition zone
Abbreviations: IZ: inhibition zone (mm); C: category; S: sensitive; R: resistant; AMP: ampicillin; CN: gentamicin; LEV: levofloxacin; VA: vancomycin; MTZ: metronidazole; MEM: meropenem; RD: rifampicin; CRO: ceftriaxone

**Table 4.** The result of sensitivity test of Gram-negative on adult sepsis patient

| Bacteria<br>(Bacteria code) | Antibiotics |  |  |  |  |  |  |  |  |  |  |  |  |  |  |  |
| --- | --- | --- | --- | --- | --- | --- | --- | --- | --- | --- | --- | --- | --- | --- | --- | --- |
|  | MEM<sup>1</sup> |  | CN<sup>1</sup> |  | LEV<sup>1</sup> |  | VA<sup>1</sup> |  | MTZ<sup>2</sup> |  | AMP<sup>2</sup> |  | RD<sup>2</sup> |  | CRO<sup>2</sup> |  |
|  | IZ | C | IZ | C | IZ | C | IZ | C | IZ | C | IZ | C | IZ | C | IZ | C |
| *Acinetobacter baumannii* (625 D) | 25,8 | S | 16 | S | 23,5 | S | - | R | - | R | 17 | R | 12,3 | R | 15 | R |
| *Escherichia coli* (646 D) | 22,6 | S | 19,75* | R | 23,25* | R | 11* | R | - | R | - | R | 14,7 | R | - | R |
| *Escherichia coli* (770 D) | 30 | S | 7,7 | R | 26,7 | S | 25* | R | - | R | - | R | 21,7 | S | 14,7 | R |
| *Escherichia coli* (866 D) | 30 | S | 19 | R | 26 | R | 21 | S | - | R | - | R | 16 | R | 12,8 | R |
| *Pseudomonas aeruginosa* (790 D) | 34 | S | 18,8 | S | 27,8 | S | - | R | - | R | - | R | 13* | R | 17,5 | S |

<sup>1</sup>: antibiotic on antibiotic guideline for sepsis patient at Dr. Moewardi regional general hospital; <sup>2</sup>: antibiotic on prescription for sepsis patient
\*: irradical; -: no inhibition zone
Abbreviations: IZ: inhibition zone (mm); C: Category; S: sensitive; R: resistant; AMP: ampicillin; CN: gentamicin; LEV: levofloxacin; VA: vancomycin; MTZ: metronidazole; MEM: meropenem; RD: rifampicin; CRO: ceftriaxone

In the result, all of Gram-negative were resistant to ampicillin and metronidazole, but sensitive against meropenem. *Acinetobacter baumannii* showed resistant to some antibiotics such as ampicillin, vancomycin, metronidazole, rifampicin, and ceftriaxone. Resistance was also shown *Pseudomonas aeruginosa* against ampicillin, vancomycin, metronidazole, and rifampicin. For *Acinetobacter baumannii* and *Pseudomonas aeruginosa* were sensitive to gentamicin, levofloxacine, and meropenem whereas *Pseudomonas aeruginosa* was sensitive to ceftriaxone.

Three of seven bacteria isolates were *Escherichia coli.* Three *Escherichia coli* were resistant against ampicillin, gantamicin, metronidazole, and ceftriaxone. But three *Escherichia coli* were sensitive to meropenem. Two of the three *Escherichia coli* showed their resistance to levofloxacine and vancomycin. On meropenem, three *Escherichia coli* were sensitive to meropenem, but occured reduction of inhibition zone, 30 mm and 22,6 mm (table 4)

### Resistance pattern of Gram-positive

29 isolates were earned from adult sepsis that consist of two isolates (primary data) and 27 isolates (secondary data) (table 2). Resistance patterns were performed to all of these isolates and the results were shown on fig. 2. Resistance also occured against erithromycin on *Staphylococcus epidermidis*, *Staphylococcus haemolyticus*, and *Staphylococcus hominis* (Fig. 2).

**Fig. 1.**
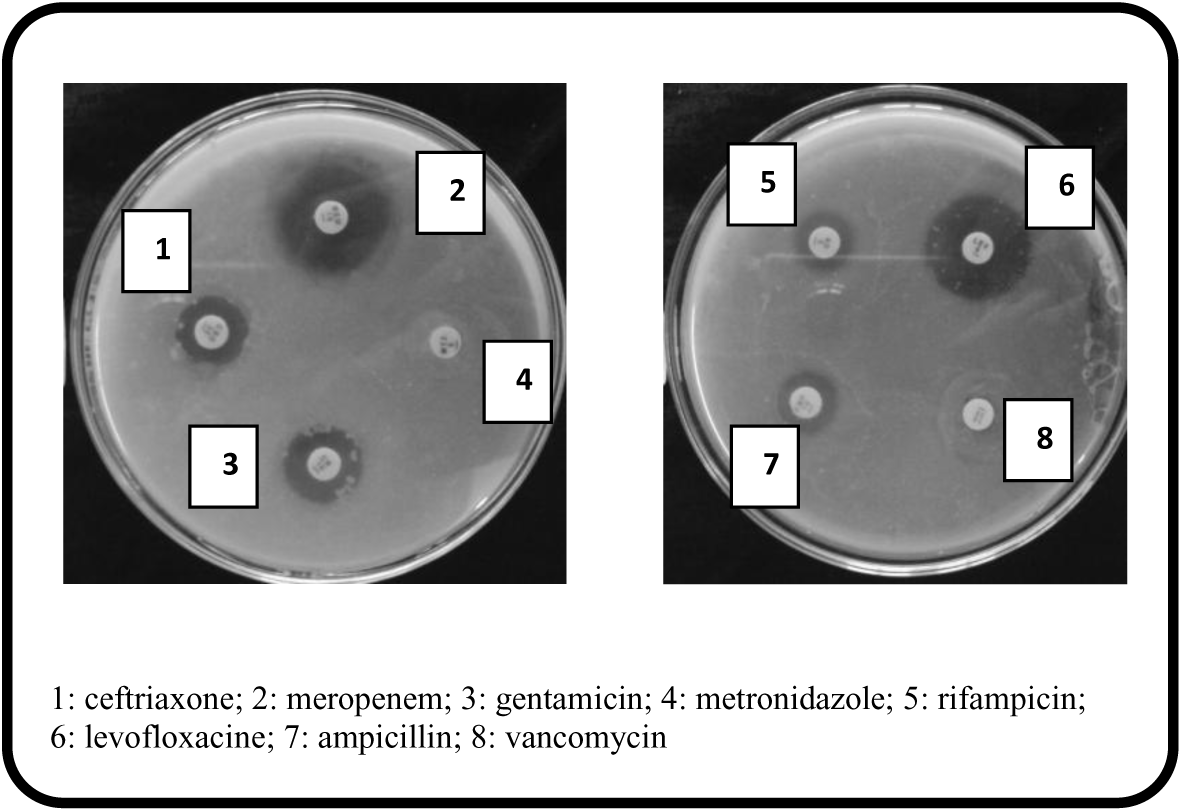
Sensitivity test result by *disc diffusion* method on *Acinetobacter baumannii*

**Fig. 3.**
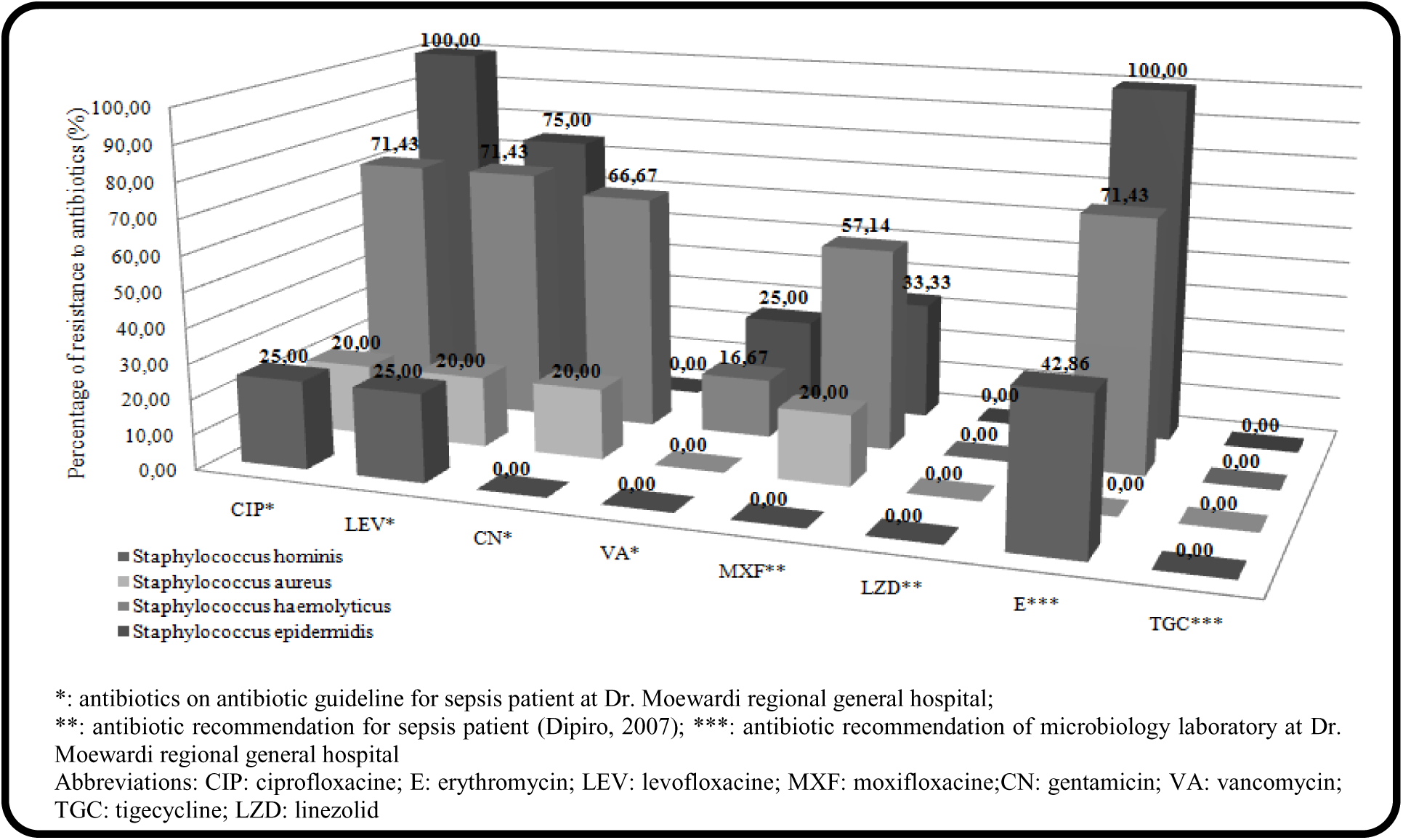
Resistance pattern of Gram-positive on adult sepsis patient at Dr. Moewardi regional general hospital in 2014 *: antibiotics on antibiotic guideline for sepsis patient at Dr. Moewardi regional general hospital;

The test result showed that there were some bacteria that resistant against antibiotics. *Staphylococcus aureus* dan *Staphylococcus haemolyticus* were resistant to vancomycin. On the other hand, *Staphylococcus hominis* was not resistant against antibiotics that were used as antibiotic recommendation for sepsis, such as gentamicin and vancomycin. All of isolated Gram-positive bacteria were sensitive against linezolide and tigecycline. *Staphylococcus aureus, Staphylococcus hominis*, and *Staphylococcus epidermidis* were sensitive to moxifloxacin, so it can be used as antibiotic for sepsis. Dipiro et al. reported that moxifloxacine and linezolid could be used as empiric antibiotic for sepsis patient. Need to be re-evaluated on antibiotic recommendation for adult sepsis patient [10].

### Resistance pattern of Gram-negative

24 Gram-negative bacteria were earned from adult sepsis patient consist of 5 bacteria isolates from primary data and 24 isolates from secondary data. *Escherichia coli* is natural bacteria in colon, but *Escherichia coli* was found on sepsis patient. Not only *Escherichia coli* but also *Acinetobacter baumannii*, *Klebsiella pneumoniae*, *Achromobacter xylosoxidant* and *Pseudomonas aeruginosa* were found on sepsis patient.

Based on the research, *Klebsiella pneumoniae*, *Acinetobacter baumannii*, and *Escherichia coli* were resistant against levofloxacine, ciprofloxacine, gentamicin, amikacin. Otherwise, all of bacteria were sensitive against meropenem. Empiric antibiotic therapy for sepsis could be used meropenem [10, 11]. Southwick reported that initial antibiotic, combination of ampicillin and amikacin, could be given when patient is diagnosed sepsis [11].

### Discussion Sepsis prevalence

Based on table 1, the state could be explained because many diseases such as influenza, pneumonia, septicemia included in the top 10 of death causes on elderly. It occurred due to the reduction of immune system [12]. A number of sepsis incidence was higher on man than woman. It was related by difference of immune responses on man and woman. Woman has good immune responses than man since woman have higher estrogen hormon. Estrogen has a role for increasing adaptive immune response. Another factor was amount of TNF (Tumor Necrosis Factor) which woman had higher than man [13].

Based on table 2, known that almost bacteria on adult sepsis was Gram-positive bacteria. Abe et al. explained that 259 sepsis patients, 65,9% were Gram-positive and 27% were Gram-negative [14]. At the same condition at Dr. Moewardi general hospital, 54,71% were Gram-positive and 45,29% Gram-negative on adult sepsis patient. It can be clarified that sepsis patient with Gramnegative bacteria had IL-6 and CRP (C-Reactive Protein) higher than Gram-positive, therefore it will make a difference mechanism of PAMPs (Pathogen-Associated Molecular Patterns) [14]. IL-6 is cytokine to regulate and increase immune system against infections and tumors [15].

### Resistance of Gram-positive

*Staphylococcus epidermidis* is one of natural bacteria on skin and mucosa. But, the bacteria sometimes causes septicemia especially people with immune system disturbance. *Staphylococcus epidermidis* had an ability to form biofilm [16]. Biofilm formation by bacteria will cause resistant against antibiotic. Resistance mechanism of biofilm occured by preventing antibiotic to reach its target and decreasing efficacy of antibiotic [17]. Ziebuhr et al. reported that *Staphylocoocus* epidermidis, isolated from blood, had a capability to form biofilm [16]. Biofilm forming will increase bacteria resistance against antibiotic and inhibit antibiotic to reach its target [18].

*Staphylococcus aureus* was bacteria that cause sepsis [4]. *Staphylococcus aureus* was resistant against gentamicin because of antibiotic modification by acetyltransferase? and phosphotransferase. On the other hand, another resistance mechanism on quinolone was the reduction of antibiotic affinity to DNA enzyme (topoisomerase IV and girase) [19]. Research on Gatermann et al. revealed that *Staphylococcus epidermidis*,

*Staphylococcus haemolyticus*, and *Staphylococcus hominis* were the top 3 bacteria that resistant against erithromycin by percentage 62,5%, 89,8%, dan 51,4%, respectively [20].

*Staphylococcus hominis* and *Staphylococcus haemolyticus* are CNS (Coagulase-Negative Staphylococci) bacteria [21]. Resistance mechanism of CNS bacteria and *Staphylococcus aureus* by movement of plasmid (contain resistant genes) from *Staphylococci* as conjugation process [22].

### Resistance of Gram-negative

Reduction of sensitivity on meropenem happened because of metallo β laktamase. *Escherichia coli* that produce the enzyme will resistant 64-folds against meropenem. This was proven by raising MIC *Escherichia coli* with metallo β laktamase (128 μg/mL) and without it (2 μg/mL) [23].

Resistance mechanism of Gram-negative bacilli through 3 resistance mechanisms those are producing enzyme to destroy antibiotic, mutation of antibiotic target, and efflux pump. A common mechanism on Gramnegative was beta lactamase production. For *Enterobacteriaceae* groups, resistance mechanism took place by destructing antibiotic through hydrolysis. The next resistance mechanism was a mutation of DNA gyrase and DNA topoisomerase IV and another resistance mechanism was a modification of antibiotic. Antibiotic modifications occured on aminoglycoside by aminoglycoside acetyltransferase, aminoglycoside O-nucleotidultransferase, and aminoglicoside O-phosphotransferase [25].

Generally, resistance mechanism of *Pseudomonas aeruginosa* against to all of antibiotic classes through reduction of cell membrane permeability and *efflux pumps* [25]. The usage of antibiotics, for instance penicillin, cephalosporine, and carbapenem, would induce the bacteria to produce beta lactamase [26]. On the other hand, resistance mechanisms of *Acinetobacter spp* were divided into 3 categories including antibiotic inactivation by enzyme, prevention of antibiotic to reach its target, and mutation of antibiotic target. Inactivation antibiotic by *Acinetobacter* through producing beta lactamase and destruction of antibiotics such as penicillin, cephalosporine, and carbapenem. Some *Acinetobacter* strains had an ability to synthesize metallo β lactamase for hydrolysis broad spectrum antibiotics included carbapenem. Another mechanism of *Acinetobacter baumannii* resistance was through *efflux pump* or *aminoglycoside-modifying enzymes* for aminoglycoside group [27].

## Conclusion

Based on the research, we could conclude that bacteria pattern on adult sepsis patient at Dr. Moewardi general hospital was *Staphylococcus haemolyticus* (15,09%), *Staphylococcus hominis* (15,09%), *Escherichia coli* (13,21%), and *Acinetobacter baumannii* (11,32%). Resistance patterns with level of resistance more than 50% as *Staphylococcus haemolyticus* (ciprofloxacine, erithromycine, levofloxacine, moxifloxacine, and gentamycin), *Escherichia coli* (gentamycin, ciprofloxacine, and levofloxacine), *Acinetobacter baumannii* (gentamycin, ciprofloxacine, levofloxacine, and ceftazidime).

## Abbreviations

IZ: inhibition zone
S: sensitive
R: resistant
AMP: ampicillin
CN: gentamicin
LEV: levofloxacine
VA: vancomycin
MTZ: metronidazole
MEM: meropenem
RD: rifampicin
CRO: ceftriaxone
CIP: ciprofloxacine
E: erythromycin
MXF: moxifloxacine
TGC: tigecycline
LZD: linezolid
CEZ: ceftazidime
AK: amikacin
Sulb-cef: sulbactam-cefoperazone
NA: Nutrient Agar
CLSI: clinical and laboratory standards institute
BHI: brain heart infusion
CFU: colony-forming units
TNF: tumor necrosis factor
IL-6: Interleukin 6
CRP: C-reactive protein
PAMPs: pathogen-associated molecular patterns
DNA: deoxyribonucleic acid
CNS: central nervous system
MIC: minimum inhibitory concentration.

## Authors’ contributions

ADM, MK, ITDK – Study concept and design. ADM – data collection and manuscript preparation. ADM, MK, ITDK – data analysis, critically revised manuscript. ADM – Analysis and interpretation of data. All authors read and approved of the final version to be published

## Acknowledgements

Microbiology lab. of Dr. Moewardi General Hospital for providing bacteria isolates. There was no external funding source for this research.

## Acknowledgement

Gratefully to Allah SWT, laboratory staffs of Microbiology Dr. Moewardi regional general hospital, and staffs of Faculty of Pharmacy, MUS.

